# Cyclase-associated protein 1 (CAP1) represses MRTF-SRF-dependent gene expression in the mouse cerebral cortex

**DOI:** 10.1101/2024.02.26.582060

**Authors:** Sharof Khudayberdiev, Kerstin Weiss, Anika Heinze, Dalila Colombaretti, Nathan Trausch, Uwe Linne, Marco B. Rust

## Abstract

Serum response factor (SRF) is a ubiquitously expressed transcription factor essential for brain development and function. SRF activity is controlled by two competing classes of coactivators, myocardin-related transcription factors (MRTF) and ternary complex factors (TCF), which introduce specificity into gene expression programs. To date, only few brain studies investigated upstream regulatory mechanisms, which mainly focused on TCF. Since an inhibitory function of monomeric actin towards MRTF-SRF signaling is well-established, we hypothesized a regulatory role for the key actin regulator ADF/cofilin. Surprisingly, ADF/cofilin was largely dispensable for neuronal MRTF-SRF activity. Instead, reporter assays combined with pharmacological and genetic approaches in isolated mouse neurons identified cyclase-associated protein 1 (CAP1) as an important regulator of this pathway. CAP1 promotes cytosolic MRTF retention and represses neuronal MRTF-SRF signaling via an actin-dependent mechanism that requires two specific protein domains. Further, deep RNA sequencing and mass spectrometry in mutant mice proved CAP1’s *in vivo* relevance for this pathway in the cerebral cortex, and led to the identification of neuronal MRTF-SRF target genes. Together, we identified CAP1 as a novel and crucial repressor of neuronal MRTF-SRF signaling.

## Introduction

The MADS box transcription factor serum response factor (SRF) induces gene expression via binding DNA at the palindromic CC[A/T]_2_A[A/T]_3_GG consensus sequence (also known as CArG box) as part of the serum response element (SRE) (Treisman, 1986; Miralles et al., 2003; Olson and Nordheim, 2010). Studies in neuronal cultures and mice implicated SRF in multiple aspects of CNS development and function, from neuron migration and differentiation to synaptic plasticity and learning (Alberti et al., 2005; Ramanan et al., 2005; Etkin et al., 2006; Knoll et al., 2006; Wickramasinghe et al., 2008; Stritt et al., 2009; Knoll, 2010; Lu and Ramanan, 2011; Kalita et al., 2012). SRF activation occurs through interaction with signal-regulated coactivators, members of either the ternary complex factor (TCF) or the myocardin-related transcription factor (MRTF) family (Olson and Nordheim, 2010; Onuh and Qiu, 2020). These coactivators compete for SRF binding, and they dictate expression of specific gene repertoires. TCF-mediated SRF activation is mainly mediated through a pathway including Ras and extracellular signal-regulated kinase (ERK), which phosphorylates TCF and thereby induces expression of a proliferative gene repertoire (Gualdrini et al., 2016). Instead, a decline in actin monomers (G-actin) activates the MRTF-SRF pathway and induces expression of among others actin and actin-binding proteins (ABP) (Miralles et al., 2003; Esnault et al., 2014), thereby providing a regulatory loop to adequately respond to cytoskeletal changes (Olson and Nordheim, 2010). The relevance of the MRTF-SRF pathway in the brain is highlighted by mutant mice lacking MRTFA and MRTFB, which phenocopied brain-specific SRF mutants (Alberti et al., 2005; Knoll, 2010; Mokalled et al., 2010; Kalita et al., 2012). Based on these findings, an inhibitory function towards neuronal SRF activity has been proposed for proteins that depolymerize actin filaments (F-actin) and, hence, increase G-actin levels. However, only few studies investigated regulatory mechanisms upstream of neuronal SRF, and these studies mainly focused on TCF (Kalita et al., 2012; Benito and Barco, 2015). Moreover, while a cytosolic MRTF retention upon G-actin binding is well established for non-neuronal cells, results in neurons were rather contradictory (Kalita et al., 2006; Wickramasinghe et al., 2008; O’Sullivan et al., 2010; Stern et al., 2013).

By exploiting cortical neurons from gene-targeted mice, the present study aimed at identifying ABP that act upstream of neuronal MRTF-SRF signaling. Because previous studies let members of the actin-depolymerizing factor (ADF)/cofilin family emerge as essential regulators of the neuronal actin cytoskeleton relevant for brain development, synapse physiology and behavior (Bellenchi et al., 2007; Hotulainen et al., 2009; Rust et al., 2010; Flynn et al., 2012; Goodson et al., 2012; Wolf et al., 2015; Zimmermann et al., 2015; Sungur et al., 2018; Schneider et al., 2021a), we were surprised to find them largely dispensable for neuronal MRTF-SRF activity. Instead, SRF activity was strongly increased upon inactivation of cyclase-associated protein 1 (CAP1), but not of its homolog CAP2. CAP1 is an ABP with largely unknown physiological functions (Rust et al., 2020; Rust and Marcello, 2022), which has been implicated in neuronal actin regulation only recently (Schneider et al., 2021b; Heinze et al., 2022). Reporter assays in isolated neurons combined with confocal microscopy as well as pharmacological and genetic approaches revealed that CAP1 promotes cytosolic MRTF retention and represses MRTF-SRF-induced gene expression via an actin-dependent mechanism. Notably, RNAseq and mass spectrometry revealed a specific upregulation of the MRTF-SRF pathway in CAP1-deficient cerebral cortex, thereby proving CAP1’s *in vivo* relevance for MRTF-SRF signaling and identifying neuronal MRTF-SRF target genes. In summary, we identified CAP1 as a crucial repressor of neuronal MRTF-SRF signaling.

## Results

### CAP1, but not ADF/cofilin or CAP2 is crucial for neuronal SRF activity

We exploited cortical neurons isolated from embryonic day 18.5 (E18.5) mice to identify upstream regulators of neuronal MRTF-SRF signaling. To assess MRTF-SRF activity, we transfected neurons after six days *in vitro* (DIV6) with 3D.AFOS construct to express firefly luciferase under control of three minimal c-Fos promotor sequences including SRF, but excluding TCF binding sites (Fig. 1A; Hill et al., 1995). As expected, natural toxins such as cytochalasin D (CytD) or jasplakinolide (Jasp), which either inhibit actin-MRTF interaction or reduce G-actin levels (Holzinger, 2009; Olson and Nordheim, 2010), robustly increased SRF activity (Fig. 1B). Instead, the actin polymerization inhibitor latrunculin B (LatB) reduced SRF activity by 42%. This reduction did not reach significance, presumably because of overall low basal SRF reporter activity in DIV10 neurons. Hence, we can induce and, if needed, inhibit MRTF-SRF activity in cortical neurons, thereby allowing us to identify upstream regulators.

**Figure 1:**
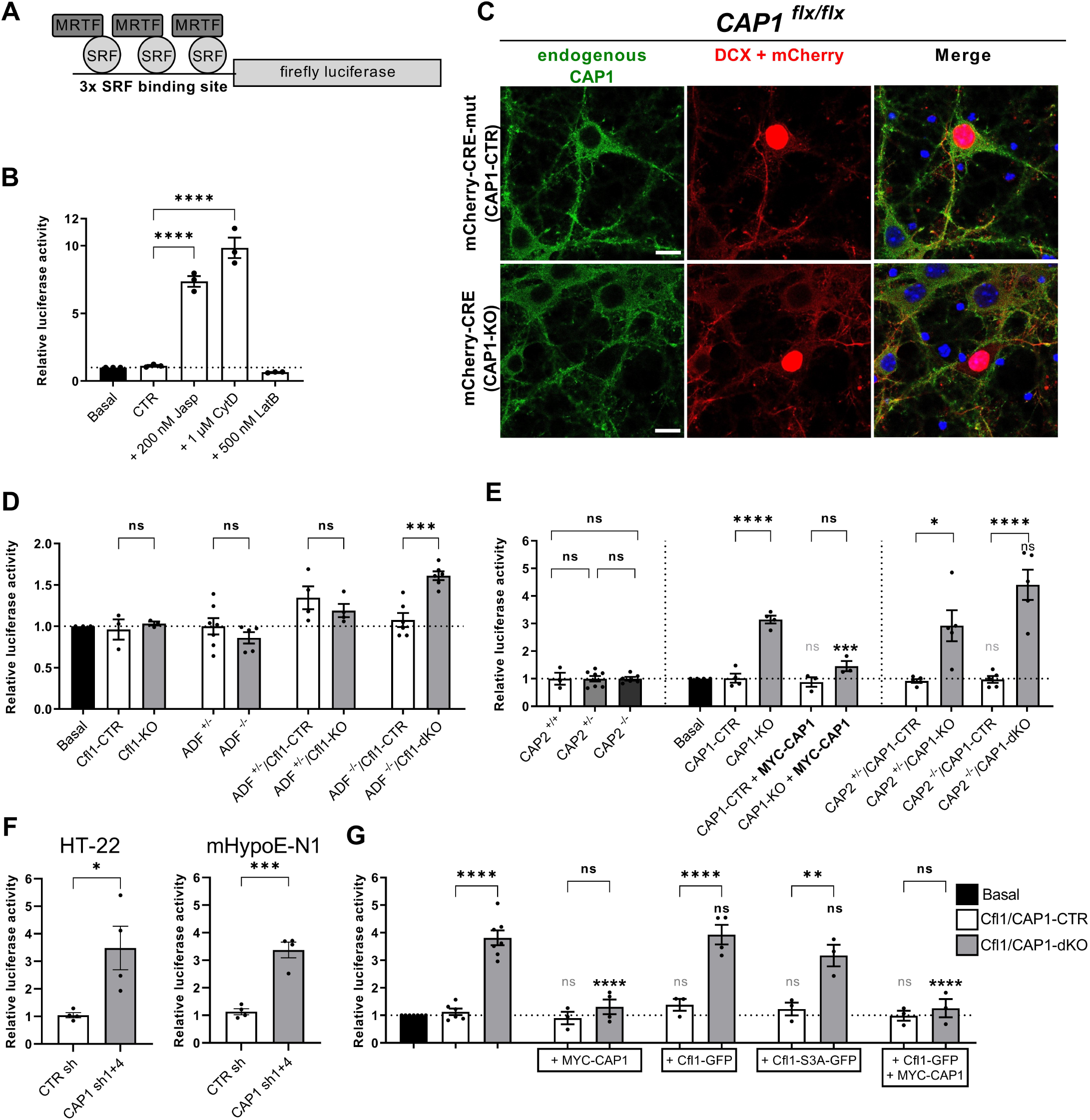
CAP1, but not ADF/cofilin or CAP2 is required for the regulation of neuronal SRF activity. **(A)** Scheme showing design of the SRF reporter 3D.AFOS that was used to determine neuronal SRF activity (Hill et al., 1995). **(B)** Graph showing increased or reduced neuronal SRF activity upon treatment with the actin drugs Jasp, CytD or LatB. Data are mean values (MV) ± standard error of the means (SEM), n=3 biological replicates (biol. rep.); one-way Anova with Dunnett’s post-hoc test. **(C)** Antibody staining showed deletion of CAP1 (green) from Cre-expressing CAP1^flx/flx^ (CAP1-KO) neurons, but not from Cre-mut-expressing CAP1^flx/flx^ (CAP1-CTR) neurons. Neurons were stained with an antibody against doublecortin (DCX, red). Cre and Cre-mut were tagged with mCherry (red). **(D)** SRF activity was unchanged in neurons lacking either ADF (ADF^-/-^) or cofilin1 (Cfl1-KO) and was only moderately increased upon inactivation of both ABP (ADF/Cfl1-dKO). Data are MV± SEM, ADF^+/-^ and ADF^-/-^ (n=5-7 biol. rep.), Cfl1-CTR and Cfl1-KO (n=3 biol. rep.), ADF/Cfl1-CTR and ADF/Cfl1-dKO (n=4-6 biol. rep.); one-way Anova with Bonferroni’s post-hoc test. **(E)** SRF activity was increased in neurons lacking CAP1 (CAP1-KO), but not CAP2 (CAP2^-/-^). SRF activity in CAP1-KO neurons was rescued by MYC-CAP1. Neurons lacking CAP1 and CAP2 (CAP1/CAP2-dKO) showed an increase in SRF activity similar to CAP1-KO neurons. Data are MV±SEM, CAP2^+/+^, CAP2^+/-^ and CAP2^-/-^ (n=3-8 biol. rep.); CAP1-CTR, CAP1-KO, CAP1-CTR + MYC-CAP1 and CAP1-KO + MYC-CAP1 (n=3-4 biol. rep.); CAP2/CAP1-CTR and CAP2/CAP1-KO (n=4 biol. rep.); one-way Anova with Sidak’s post-hoc test. **(F)** shRNA-mediated CAP1 inactivation increased SRF activity in the neuronal cell lines HT-22 and mHypoE-N1. Data are MV±SEM, n=4 biol. rep.; two-tailed Student’s t-test. **(G)** Neurons lacking cofilin1 and CAP1 (Cfl1/CAP1-dKO) showed a similar increase in SRF activity as CAP1-KO neurons. In Cfl1/CAP1-dKO neurons, SRF activity was normalized by MYC-CAP1, but not by Cfl1-GFP or a constitutively active cofilin1 mutant (Cfl1-S3A). Data are MV±SEM, n=3-7 biol. rep.; one-way Anova with Sidak’s post-hoc test. Scale bar (µm): 10 (C). ns: P≥0.05, *: P<0.05, **: P<0.01, ***: P<0.001, ****: P<0.0001. The column bars containing light grey and dark color statistical significance labels without brackets represent comparison to CTR and KO conditions, respectively

Studies from us and others identified cofilin1 and ADF as key actin regulators in neurons (Rust, 2015a, b), whose inactivation shifted the actin equilibrium towards F-actin (Rust et al., 2010; Flynn et al., 2012; Wolf et al., 2015). Further, we recently demonstrated an interaction of cofilin1 with both CAP2 and CAP1 in neurons (Pelucchi et al., 2020; Schneider et al., 2021b; Heinze et al., 2022). We therefore expected increased SRF activity in neurons lacking i) cofilin1, ii) cofilin1 and ADF, iii) CAP2 and/or iv) CAP1. To test this, we transfected DIV6 neurons isolated from either Cfl1^flx/flx^ mice, ADF^-/-^/Cfl1^flx/flx^ mice, CAP1^flx/flx^ or CAP1^flx/flx^/CAP2^-/-^ mice with the SRF reporter together with either catalytically active mCherry-Cre (Cre) or a catalytically inactive, mutant mCherry-Cre variant (Cre-mut). Immunocytochemistry demonstrated absence of CAP1 from Cre-transfected CAP1^flx/flx^ neurons (CAP1-KO), but presence in Cre-mut-transfected CAP1^flx/flx^ neurons (CAP1-CTR), thereby validating suitability of our experimental approach (Fig. 1C).

Unexpectedly, SRF activity was not different between Cfl1^flx/flx^ neurons expressing either Cre (Cfl1-KO) or Cre-mut (Cfl1-CTR; Fig. 1D). Further, it was neither increased in ADF^-/-^ neurons nor in ADF^+/-^/Cfl1-KO neurons. When compared to Cre-mut-expressing ADF^-/-^/Cfl1^flx/flx^ neurons (ADF/Cfl1-CTR), SRF activity was increased by 60% in Cre-expressing ADF^-/-^/Cfl1^flx/flx^ neurons (ADF/Cfl1-dKO; Fig. 1D). Together, only compound inactivation of cofilin1 and ADF increased SRF activity, and this increase was rather moderate.

Further, SRF activity was not different between neurons isolated from CAP2^+/+^ (CAP2-CTR), CAP2^+/-^ or CAP2^-/-^ (CAP2-KO) mice (Fig. 1E). Instead, SRF activity was increased more than threefold in CAP1-KO neurons, and it was normalized to CAP1-CTR levels by expression of MYC-, GFP- or mCherry-tagged CAP1 (Figs. 1E, S1A, S2). Similarly, shRNA-mediated CAP1 silencing increased SRF activity roughly threefold in the mouse neuronal cell lines HT-22 and mHypoE-N1 (Figs. 1F, S1B-G). When compared to Cre-mut-expressing CAP2^+/-^/CAP1^flx/flx^ neurons (CAP2/CAP1-CTR), SRF activity was increased in Cre-expressing CAP2^-/-^/CAP1^flx/flx^ neurons (CAP2/CAP1-dKO). However, the increase was only slightly higher when compared to CAP1-KO neurons, and this comparison did not reach significance (Fig. 1E). These data suggested a crucial role for CAP1, but not for CAP2 in regulating neuronal SRF activity. Since previous studies revealed a cooperation of cofilin1 and CAP1 in actin subunit dissociation and in neuron morphology (Kotila et al., 2019; Shekhar et al., 2019; Schneider et al., 2021b; Heinze et al., 2022), we tested whether compound inactivation of both ABP further increased SRF activity. Interestingly, Cre-expressing Cfl1^flx/flx^/CAP1^flx/flx^ neurons (Cfl1/CAP1-dKO) displayed an increase in SRF reporter activity very similar to CAP1-KO neurons (Fig. 1G). Notably, expression of MYC-CAP1, but neither of GFP-tagged wildtype (WT) cofilin1 (Cfl1-GFP) nor of constitutive active cofilin1 (Cfl1-S3A-GFP) was sufficient to normalize SRF activity in these neurons, thereby confirming our above-drawn conclusions that CAP1, but not cofilin1 was crucial for neuronal SRF activity.

**Figure 2:**
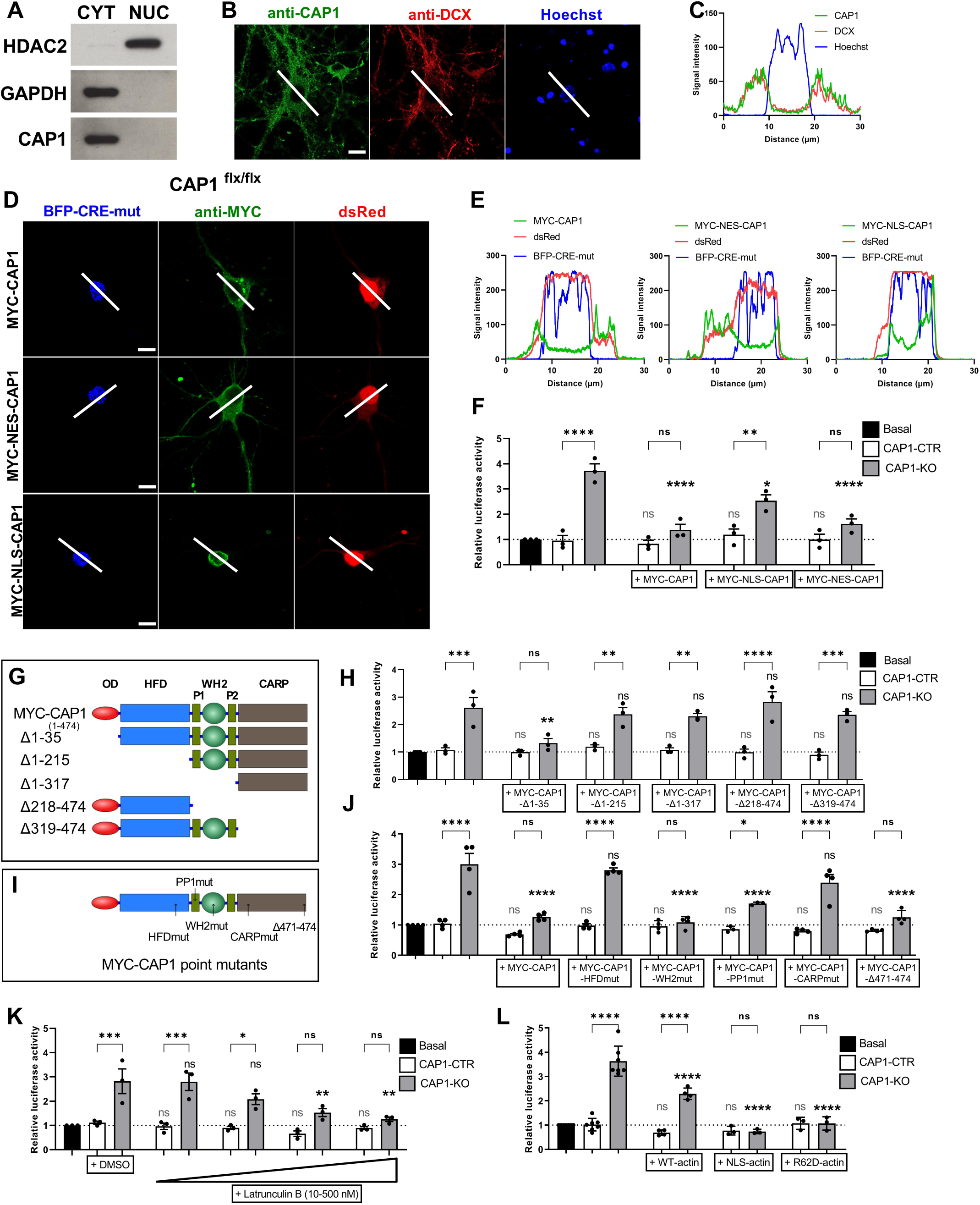
Cytosolic CAP1 controls neuronal SRF activity via an actin-dependent mechanism that requires its HFD and CARP domain. **(A)** Immunoblots of cytosol- and nucleus-enriched protein fractions from cerebral cortex from postnatal day 12 mice showing abundance of CAP1 in cytosolic fraction. GAPDH and HDAC2 proteins were detected as markers for cytosol and nucleus, respectively, to prove separation of these subcellular compartments by fractionation. **(B)** Antibody staining in DIV10 cortical neurons revealed cytosolic localization of endogenous CAP1 (green). Neurons were counterstained with an antibody against DCX (red) and the DNA dye Hoechst (blue). **(C)** Fluorescence intensity profiles for CAP1, DCX and Hoechst along white line in Fig. 1B. **(D)** Myc antibody staining (green) in DIV10 cortical neurons expressing MYC-CAP1 variants together with blue fluorescent protein (BFP)-tagged Cre-mut (blue) and *Discosoma* red fluorescent protein (dsRed, red), which were used as nuclear and volume marker, respectively. **(E)** Fluorescence intensity profiles for MYC-CAP1 variants, BFP-Cre-mut and dsRed along white line in Fig. 1D. **(F)** MYC-CAP1 or MYC-NES-CAP1 normalized SRF activity in CAP1-KO neurons. MYC-NLS-CAP1 reduced SRF activity in CAP1-KO neurons, which was still higher when compared to CAP1-CTR neurons. Data are MV±SEM, n=3 biol. rep.; one-way Anova with Sidak’s post-hoc test. **(G)** Scheme showing protein domains of full length CAP1 as well as CAP1 deletion constructs. **(H)** While MYC-CAP1-Δ1-35 normalized SRF activity in CAP1-KO neurons, none of the other tested deletion mutants reduced SRF activity in CAP1-KO neurons. Data are MV±SEM, n=3 biol. rep.; one-way Anova with Sidak’s post-hoc test. **(I)** Scheme showing CAP1 point mutations. **(J)** Similar to MYC-CAP1, CAP1-WH2mut and CAP1-Δ471-474 normalized SRF activity in CAP1-KO neurons. Instead, CAP1-PP1mut reduced SRF activity in CAP1-KO neurons, which was still higher when compared to CAP1-CTR neurons, and CAP1-HFDmut and CAP1-CARPmut did not reduce SRF activity in CAP1-KO neurons. Data are MV±SEM, n=3 biol. rep.; one-way Anova with Sidak’s post-hoc test. **(K)** The actin polymerization inhibitor LatB normalized SRF activity in CAP1-KO neurons. Data are MV±SEM, n=3 biol. rep.; one-way Anova with Sidak’s post-hoc test. **(L)** Overexpression of WT-actin reduced SRF activity in CAP1-KO neurons, which was still higher when compared to CAP1-CTR neurons. Instead, a polymerization impaired actin mutant (R62D-actin) as well as nuclear-targeted actin (NLS-actin) normalized SRF activity in CAP1-KO neurons. Data are MV±SEM, n=3 biol. rep.; one-way Anova with Sidak’s post-hoc test. Scale bars (µm): 10 (B, D). ns: P≥0.05, *: P<0.05, **: P<0.01, ***: P<0.001, ****: P<0.0001. The column bars containing light grey and dark color statistical significance labels without brackets represent comparison to CTR and KO conditions, respectively.

### Cytosolic CAP1 controls SRF activity via an actin-dependent mechanism, which requires its HFD and CARP domain

To unravel how CAP1 controls SRF activity, we first determined its subcellular localization. Immunoblots of cerebral cortex lysates from postnatal day 12 mice revealed CAP1 enrichment in cytosolic and absent from nuclear fraction (Fig. 2A). Antibody staining against endogenous CAP1 or MYC in MYC-CAP1-expressing neurons confirmed cytosolic CAP1 localization in neurons (Fig. 2B-E), thereby suggesting that cytosolic CAP1 controls SRF activity. However, a previous study reported cytosolic and nuclear localization for CAP2 and suggested that it acts as dual compartment protein (Peche et al., 2007). We therefore determined SRF activity in CAP1-KO neurons upon expression of CAP1 variants carrying either a nuclear localization signal (MYC-NLS-CAP1) or a nuclear export signal (MYC-NES-CAP1), which showed the expected abundance in the nucleus or cytosol, respectively (Fig. 2D-E). MYC-NES-CAP1 normalized SRF activity in CAP1-KO neurons, very similar to MYC-CAP1 (Fig. 2F). Instead, MYC-NLS-CAP1 did not normalize SRF activity in CAP1-KO neurons, albeit it was reduced when compared to CAP1-KO neurons. Hence, CAP1 activity in the cytosol, but not in the nucleus, was sufficient for regulating neuronal SRF activity.

Next, we determined SRF activity in CAP1-KO upon expression of CAP1 deletion constructs to identify protein domain relevant for SRF regulation (Fig. 2G). A deletion mutant (CAP1-Δ1-35) lacking the N-terminal 35 amino acid (AA) residues including the oligomerization domain (OD) as well as two Arg-Leu-Glu (RLE) repeats normalized SRF activity in CAP1-KO neurons (Fig. 2H). Instead, CAP1 mutants lacking the entire N-terminal part (CAP1-Δ1-215) or the N-terminal part together with the central region (CAP1-Δ1-317) were not able to rescue reporter activity, similar to CAP1 variants lacking either the central region together with the C-terminal part (CAP1-Δ218-474) or the C-terminal part only (CAP1-Δ319-474). Hence, CAP1’s OD and RLE repeats were dispensable for SRF regulation, while protein domains of the N- as well as the C-terminal part were both relevant.

The N-terminal part contains a helical folded domain (HFD), while the C-terminal part mainly consists of a CARP domain (Ono, 2013; Rust et al., 2020; Rust and Marcello, 2022). In order to pinpoint CAP1’s domains relevant for SRF regulation, we determined SRF activity in CAP1-KO neurons upon expression of mutant CAP1 variants that reportedly 1) diminished its interaction with actin subunits at F-actin pointed-end (CAP1-HFDmut), 2) disturbed binding to its poly-proline domain 1 (CAP1-PP1mut) or impaired its nucleotide exchange activity on G-actin by mutating either 3) the central WH2 domain (CAP1-WH2mut), 4) the CARP domain (CAP1-CARPmut) or 5) deleting the C-terminal four AA residues (CAP1-Δ471-474) (Fig. 2I; Kotila et al., 2018; Kotila et al., 2019). While CAP1-WH2mut, CAP1-PP1mut and CAP1-Δ471-474 normalized SRF activity in CAP1-KO neurons, CAP1-HFDmut and CAP1-CARPmut both failed in reducing SRF activity (Figs. 2J, S2). Outcome was independent from the tagged proteins (MYC and mCherry) or their size relative to CAP1. Hence, HFD and CARP domain, which both have been implicated in actin dynamics (Kotila et al., 2018; Kotila et al., 2019), were relevant for regulating neuronal SRF activity, suggesting that CAP1 controls SRF via an actin-dependent mechanism.

Indeed, SRF activity in CAP1-KO neurons was normalized by the actin polymerization inhibitor LatB (Fig. 2K). Upon treatment with 500 nM LatB, SRF activity was not different between CAP1-CTR and CAP1-KO neurons. Moreover, increasing cellular G-actin levels by expressing either WT-actin or an actin mutant (R62D-actin) impaired in polymerization and CAP1 binding, reduced SRF activity in CAP1-KO neurons (Fig. 2L). R62D-actin expression fully rescued SRF activity in CAP1-KO neurons, similar to NLS-actin, which likewise can’t interact with CAP1 due to its nuclear localization (McCormack et al., 2001). Together, our data revealed that neuronal SRF activity is controlled by cytosolic CAP1 via an actin-dependent mechanism.

### CAP1 represses neuronal SRF activity via promoting cytosolic MRTF retention

Since MRTF is able to translate cytosolic actin changes into nuclear SRF activity, we assumed that CAP1 controls MRTF activity (Olson and Nordheim, 2010). Indeed, inhibiting MRTF activity by exploiting the small-molecule inhibitor CCG-203971 or by shRNA-mediated knockdown normalized SRF activity in CAP1-KO neurons (Fig. 3A-B). Further, we overexpressed either Flag-MRTFA or MYC-MRTFB in cortical neurons, which both increased SRF activity in CAP1-KO neurons, but not in CAP1-CTR neurons (Fig. 3C-D). Expression of MYC-CAP1 overcame these effects and reduced SRF activity in CAP1-KO neurons overexpressing either MRTFA or -B to CAP1-CTR levels. Instead, ΔN-MRTFA, a mutant variant lacking the inhibitory actin-binding RPEL domain (Miralles et al., 2003), strongly increased SRF activity in CAP1-CTR neurons, but only moderately in CAP1-KO neurons, and MYC-CAP1 failed to reduce SRF activity in ΔN-MRTFA-expressing neurons (Fig. 3E). Hence, our data revealed increased MRTF activity in CAP1-KO neurons and that MRTF regulation by CAP1 required MRTF’s RPEL domain.

**Figure 3:**
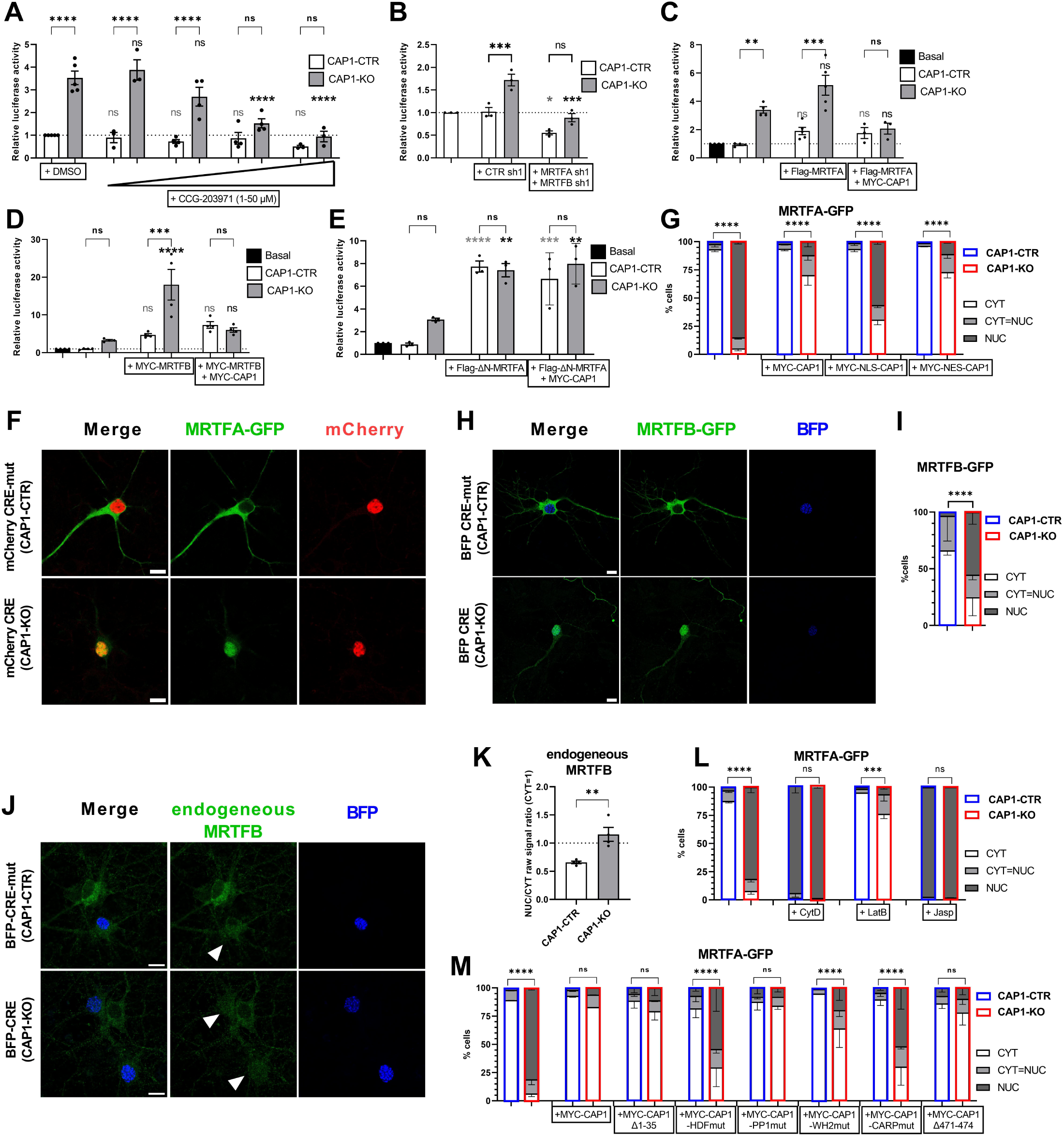
Nucleo-cytosolic shuttling of MRTF links cytosolic CAP1 activity to neuronal SRF activity. **(A)** The small-molecular inhibitor CCG-203971 as well as **(B)** shRNA-mediated inactivation of MRTFA and MRTFB normalized SRF activity in CAP1-KO neurons. Data are MV±SEM, n=3 biol. rep.; one-way Anova with Sidak’s post-hoc test. Overexpression of **(C)** MRTFA or **(D)** MRTFB increased SRF activity in CAP1-KO neurons, but neither in MYC-CAP1-expressing CAP1-KO neurons nor in CAP1-CTR neurons. Data are MV±SEM, n=3-5 biol. rep.; one-way Anova with Sidak’s post-hoc test. **(E)** Instead, overexpression of a mutant MRTFA variant lacking the actin-binding RPEL domain (ΔN-MRTFA) increased SRF activity in CAP1-CTR neurons, but not in CAP1-KO neurons either expressing MYC-CAP1 or not. Data are MV±SEM, n=4 biol. rep.; one-way Anova with Sidak’s post-hoc test. **(F)** Micrographs of CAP1-CTR and CAP1-KO neurons expressing MRTFA-GFP (green). mCherry-tagged Cre or Cre-mut (red) was used as nuclear marker. **(G)** Fractions of CAP1-CTR and CAP1-KO neurons expressing or not MYC-CAP1, MYC-NLS-CAP1 or MYC-NES-CAP1 displaying MRTFA-GFP localization in cytosol (CYT), nucleus (NUC) or equal distribution between both compartments (CYT=NUC). For each experimental group within a biol. rep., 114-506 neurons were analyzed. Results were converted to fractions (percentage) and MV±SEM for n=2-3 biol. rep. was calculated; χ^2^ test. **(H)** Micrographs of CAP1-CTR and CAP1-KO neurons expressing MRTFB-GFP (green). BFP-tagged Cre or Cre-mut (blue) was used as nuclear marker. **(I)** Fractions of CAP1-CTR and CAP1-KO neurons displaying MRTFB-GFP localization in CYT, NUC or both compartments (CYT=NUC). For each experimental group within a biol. rep., 83-136 neurons were analyzed. Results were converted to fractions (percentage) and MV±SEM for n=3 biol. rep. was calculated; χ^2^ test. **(J)** Antibody staining against endogenous MRTFB in CAP1-CTR and CAP1-KO neurons. BFP-Cre or -Cre-mut (blue) was used as nuclear marker. **(K)** NUC/CYT ratio of MRTFB immunoreactivity was increased in CAP1-KO neurons. Data are MV±SEM, n=4 biol. rep.; two-tailed Student’s t-test. **(L)** LatB treatment normalized MRTFA-GFP localization in CAP1-KO neurons, while CytD and Jasp shifted almost all MRTFA-GFP into the nucleus of CAP1-CTR and CAP1-KO neurons. For each experimental group within a biol. rep., 62-416 neurons were analyzed. Results were converted to fractions (percentage) and MV±SEM for n=3 biol. rep. was calculated; χ^2^ test. **(M)** MYC-CAP1-Δ1-35, MYC-CAP1-PP1mut and MYC-CAP1-Δ471-474 normalized MRTFA-GFP localization in CAP1-KO neurons. MYC-CAP1-WH2 mutant increased CYT fraction, but did not fully rescue MRTFA-GFP localization in CAP1-KO neurons. Instead, MYC-CAP1-HFDmut and MYC-CAP1-CARPmut failed in altering MRTFA-GFP localization in CAP1-KO neurons. For each experimental group within a biol. rep., 76-243 neurons were analyzed. Results were converted to fractions (percentage) and MV±SEM for n=3 biol. rep. was calculated; χ^2^ test. Scale bars (µm): 10 (F, H, J). ns: P≥0.05, *: P<0.05, **: P<0.01, ***: P<0.001, ****: P<0.0001. The column bars containing light grey and dark color statistical significance labels without brackets represent comparison to CTR and KO conditions, respectively.

Nucleo-cytosolic MRTF shuttling emerged as an important mechanism of SRF regulation in various cell types (Olson and Nordheim, 2010). However, findings in neurons were contradictory as some studies showed exclusive nuclear MRTF localization (Kalita et al., 2006; Wickramasinghe et al., 2008; O’Sullivan et al., 2010; Stern et al., 2013), which would be difficult to link to a regulation by cytosolic CAP1. We therefore determined MRTF localization in cortical neurons to test whether it depended on CAP1. To do so, we expressed either GFP-tagged MRTFA or -B and calculated neuron fractions with cytosolic (CYT), nuclear (NUC) or equal distribution (CYT=NUC) between both compartments (Fig. S3). The vast majority of CAP1-CTR neurons expressing MRTFA-GFP or MRTFB-GFP belonged to the CYT fraction and only a few displayed predominant NUC expression (Fig. 3F-I). Conversely, almost all MRTFA-GFP-expressing CAP1-KO neurons and the majority of MRTFB-GFP-expressing CAP1-KO neurons belonged to the NUC fraction. By antibody staining against endogenous MRTFB we confirmed increased nuclear levels in CAP1-KO neurons, in which the NUC/CYT ratio of MRTFB immunoreactivity was twice as high as in CAP1-CTR neurons (Fig. 3J-K). Together, CAP1 inactivation shifted MRTF localization towards the nucleus.

Next, by using MRTFA-GFP localization as a readout, we determined how CAP1 controls subcellular MRTF distribution. Apart from CAP1-KO neurons, we included ADF/Cfl1-dKO neurons in these experiments, in which moderately increased SRF activity was associated with a moderate shift towards nuclear localized MRTFA-GFP (Fig. S3C). MRTFA localization in CAP1-KO and ADF/Cfl1-dKO neurons was normalized by LatB, while treatment with either CytD or Jasp shifted almost all MRTFA into the nucleus independent of the presence of CAP1 or ADF/cofilin (Figs. 3L, S3B). Hence, manipulating the actin cytoskeleton robustly affected MRTF localization, thereby challenging earlier studies (Kalita et al., 2006).

Expression of either MYC-CAP1 or MYC-NES-CAP1 in CAP1-KO neurons shifted MRTFA towards the cytosol (Figs. 3G, S3C). Instead, MYC-NLS-CAP1 only partially rescued MRTFA localization, and most CAP1-KO neurons still belonged to NUC fraction. Moreover, expression of CAP1-Δ1-35, CAP1-PP1mut or CAP1-Δ471-474 normalized MRTFA localization in CAP1-KO neurons, and CAP1-WH2mut strongly decreased NUC and CYT=NUC fractions (Fig. 3M). Instead, expression of CAP1-HFDmut or CAP1-CARPmut both failed in altering MRTFA localization in CAP1-KO neurons. These results were in good agreement with our SRF reporter assay, in which MYC-CAP1-HFDmut and MYC-CAP1-CARPmut failed in reducing SRF activity in CAP1-KO neurons. Taken together, our data presented up to here revealed that cytosolic CAP1 controls MRTF localization and SRF activity via an actin-dependent mechanism, which requires MRTF’s RPEL domain as well as CAP1’s HFD and CARP domain, but not other CAP1 domains such as OD, RLE repeats, PP1 or WH2.

### CAP1 inactivation in mouse brain specifically increased MRTF-SRF signaling

Having established CAP1 as a repressor MRTF-SRF activity in cortical neurons, we next tested whether CAP1 exerted a similar function *in vivo*. We therefore performed deep RNA sequencing (RNAseq) on cerebral cortex lysates from E18.5 brain-specific CAP1-KO mice (CAP1^flx/flx,Nestin-Cre^) and CAP1^flx/flx^ (CTR) littermates (Schneider et al., 2021b). We identified in total 15,942 genes of which 154 genes were differentially expressed (FDR<0.05) between both groups (Fig. 4A-B, Tab. S1). In CAP1-KO mice, 90 genes were upregulated and 64 genes including CAP1 were downregulated (Tab. S2). Next, we performed gene ontology (GO) analyses focusing on the domains ‘biological process’ (GOBP) and ‘cellular component’ (GOCC). GOBP analysis revealed an enrichment of genes implicated in sterol biosynthesis and metabolism among downregulated genes (Fig. 4C). Conversely, genes implicated in cytoskeleton organization, actomyosin contractility or F-actin-based processes were found among upregulated genes. Accordingly, GOCC analysis revealed location of upregulated genes at cellular compartments enriched in cytoskeletal elements such as focal adhesions or adherens junctions. Kyoto Encyclopedia of Genes and Genomes (KEGG) analysis revealed an enrichment of genes that have been associated with dilated cardiomyopathy (DCM) and hypertrophic cardiomyopathy (HCM) among genes upregulated in CAP1-KO mice, while genes associated with terpenoid biosynthesis were enriched among downregulated genes. Notably, a priori gene set enrichment analysis (GSEA) revealed an enrichment of SRF targets among those genes upregulated in CAP1-KO mice.

**Figure 4:**
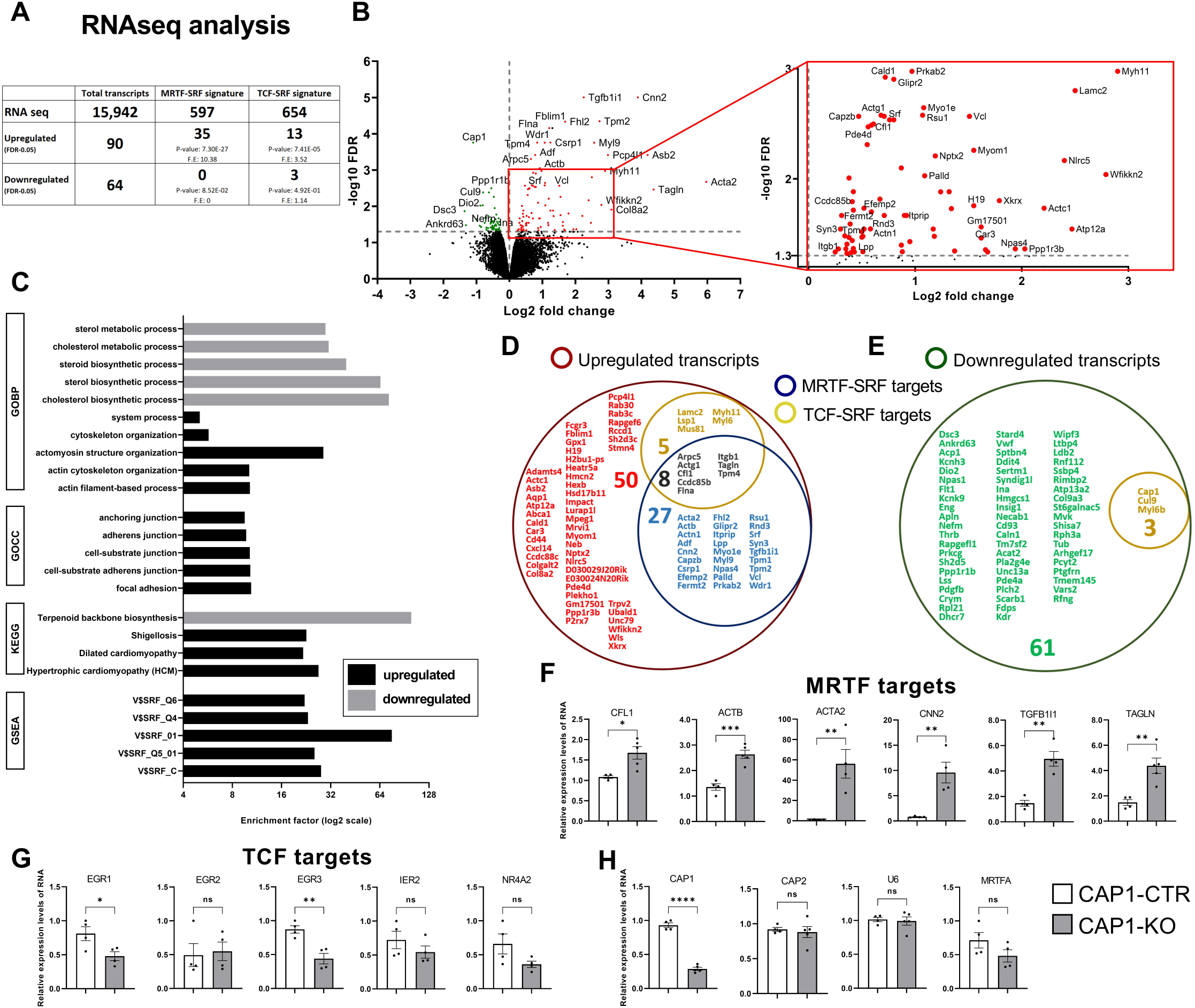
RNAseq revealed a specific upregulation of MRTF-SRF target genes in CAP1-KO brain. **(A)** Table showing total transcript number as well as MRTF-SRF and TCF-SRF targets identified in RNAseq analysis of E18.5 cerebral cortex. Moreover, table includes transcript numbers either up- or downregulated in CAP1-KO cerebral cortex. Statistical analysis for enrichment was performed with hypergeometric test. MRTF-SRF and TCF-SRF signature genes were obtained from previous studies (Esnault et al., 2014; Gualdrini et al., 2016). F.E.: fold enrichment. **(B)** Volcano plot showing transcripts downregulated (green) or upregulated (red) in E18.5 CAP1-KO cerebral cortex. Red box indicates part of the volcano plot shown at higher magnification. **(C)** Gene ontology (GO) analyses focusing on the domains ‘biological process’ (GOBP) and ‘cellular component’ (GOCC), Kyoto Encyclopedia of Genes and Genomes (KEGG) analysis as well as gene set enrichment analysis (GSEA) of transcripts dysregulated in E18.5 CAP1-KO cerebral cortex. Bubble chart separating transcripts upregulated **(D)** and downregulated **(E)** in E18.5 CAP1-KO cerebral cortex into groups either regulated by MRTF-SRF only (blue), by TCF-SRF only (yellow), by both pathways (black) or neither by MRTF-SRF or TCF-SRF (red: upregulated, green: downregulated). qPCR with cerebral cortex lysates from CAP1-KO mice **(F)** confirmed upregulation of selected MRTF-SRF targets, **(G)** revealed reduced or unchanged expression of TCF-SRF-specific targets, and **(H)** showed expression levels of CAP1, CAP2, U6 and MRTFA. Data are MV±SEM, n=4-5 biol. rep.; two-tailed Student’s t-test. ns: P≥0.05, *: P<0.05, **: P<0.01, ***: P<0.001, ****: P<0.0001.

The latter finding forced us to compare genes dysregulated in CAP1-KO mice to recent studies that comprehensively characterized SRF target gene sets (Esnault et al., 2014; Gualdrini et al., 2016). This comparison confirmed 40 established SRF targets among 90 upregulated genes in CAP1-KO cerebral cortex (44.4%), while only 3 SRF targets (4.7%) were found among 64 downregulated genes (Fig. 4D-E; Tab. S2). The coactivators MRTF and TCF compete for SRF’s DNA-binding domain and induce expression of specific, but sometimes overlapping gene sets (Esnault et al., 2014; Gualdrini et al., 2016). We therefore sub-grouped dysregulated SRF targets into MRTF-SRF and TCF-SRF gene sets. Of 40 SRF targets upregulated in CAP1-KO mice, only 5 genes belonged to TCF-SRF group, while 27 genes belonged to MRTF-SRF group. Expression of the remaining 8 genes is believed to be controlled by both pathways (Fig. 4D). Conversely, all downregulated SRF targets (3 genes) in CAP1-KO mice belonged to the TCF-SRF group (Fig. 4E). Together, our analyses revealed 7.37-fold enrichment of MRTF-SRF target genes among upregulated genes in CAP1-KO mice, while TCF-SRF target genes were either less affected or downregulated. These findings were in line with our data in cortical neurons, in which CAP1 controls SRF via MRTF.

Next, we performed quantitative PCR (qPCR) on cerebral cortex lysates to validate RNAseq data. qPCR confirmed upregulation of selected MRTF-SRF targets such as Acta2, Actb, Cfl1, Cnn2, Tagln and Tgfb1i1 in CAP1-KO mice (Fig. 4F), while expression of genes selectively controlled by TCF-SRF were either downregulated (Egr1, Egr3) or unaltered (Egr2, Ier2, Nr4a2; Fig. 4G). Further, qPCR confirmed reduced CAP1 expression as well as unaltered expression of CAP2, U6 or MRTFA in CAP1-KO mice (Fig. 4H). Hence, RNA seq revealed a specific upregulation of MRTF-SRF target genes in cerebral cortex of CAP1-KO mice.

### Mass spectrometry confirmed upregulation of the MRTF-SRF pathway

Next, we performed mass spectrometry on cerebral cortex lysates to test whether changes on mRNA level in CAP1-KO mice were present on protein level, too. We identified in total 2,545 proteins, of which 65 were differentially expressed (FDR<0.05) between both groups (Fig. 5A-B; Tab. S3). In CAP1-KO mice, 37 proteins were upregulated and 28 downregulated (Fig. 5A, Tab. S4). As expected, CAP1 showed strongest downregulation among all proteins (Fig. 5B). GOCC analysis revealed an enrichment of proteins associated with neuro-/intermediate filaments among proteins downregulated in CAP1-KO mice, while proteins associated with cytoskeletal elements (focal adhesions, cell and anchoring junctions, contractile fibers) were enriched among upregulated proteins (Fig. 5C). Notably, a priori GSEA revealed an enrichment of SRF targets among upregulated proteins (Fig. 5C). More detailed analysis of differentially expressed proteins revealed an enrichment (4.25-fold) of MRTF-SRF targets, but not of TCF-SRF targets among upregulated proteins (Fig. 5A, D). Conversely, neither MRTF-SRF nor TCF-SRF targets were enriched among 28 downregulated proteins (Fig. 5A, E). Hence, mass spectrometry data largely confirmed a dysregulation of the MRTF-SRF pathway in CAP1-KO mice. Immunoblot analyses confirmed mass spectrometry data by demonstrating increased expression levels for vinculin, twinfilin1 and cofilin1 in CAP1-KO mice, although the latter missed significance by margin (Fig. 5F-G). Further, they confirmed unaltered protein levels for ADF and GAPDH in CAP1-KO lysates.

**Figure 5:**
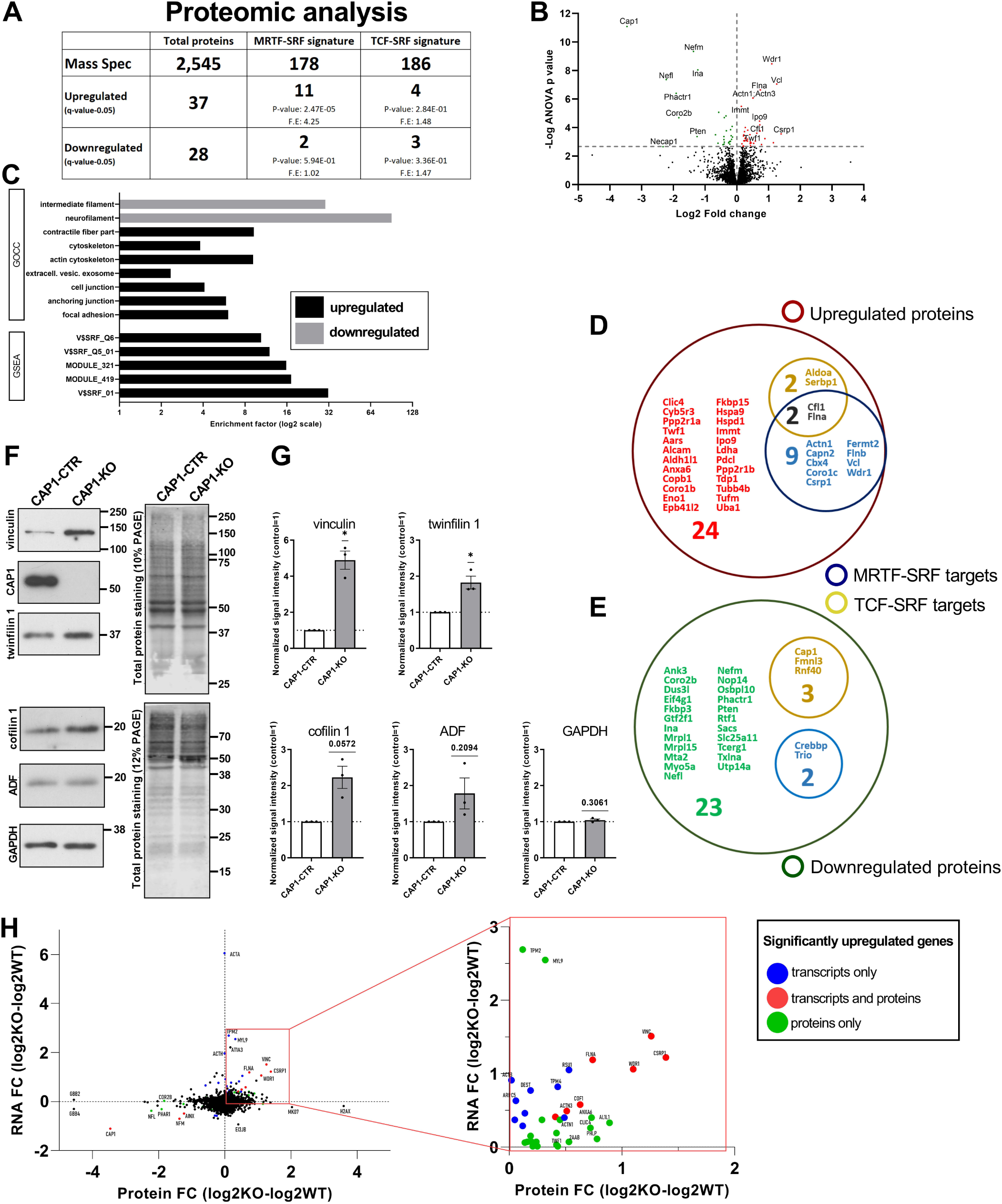
CAP1-KO brains show upregulated expression of MRTF-SRF targets. **(A)** Table showing total protein number as well as MRTF-SRF and TCF-SRF targets identified by mass spectrometry in E18.5 cerebral cortex. Moreover, table includes protein numbers either up- or downregulated in CAP1-KO cerebral cortex. Statistical analysis for enrichment was performed with hypergeometric test. MRTF-SRF and TCF-SRF signature genes were obtained from previous studies (Esnault et al., 2014; Gualdrini et al., 2016). F.E.: fold enrichment. **(B)** Volcano plot showing proteins downregulated (green) or upregulated (red) in E18.5 CAP1-KO cerebral cortex. **(C)** GOBP and GOCC analysis of proteins dysregulated in E18.5 CAP1-KO cerebral cortex. Bubble charts separating proteins either **(D)** upregulated or **(E)** downregulated in E18.5 CAP1-KO cerebral cortex into groups either regulated by MRTF-SRF only (blue), by TCF-SRF only (yellow), by both pathways (black) or neither by MRTF-SRF or TCF-SRF (red: upregulated, green: downregulated). **(F)** Immunoblots of E18.5 cerebral cortex lysates with antibodies against selected proteins. GADPH was used as loading control. Additionally, total protein stain confirmed equal loading. **(G)** Quantification of immunoblots revealed increased expression levels for vinculin, twinfilin1 and cofilin1 in E18.5 CAP1-KO cerebral cortex, although the latter missed significance by margin. Instead, expression levels of ADF was unchanged. Immunoblots thereby confirmed mass spectrometry data. Data are MV±SEM, n=3 biol. rep.; two-tailed Student’s t-test. **(H)** Comparison of RNAseq and mass spectrometry data by plotting fold changes of transcripts against corresponding proteins. Color code indicate transcripts changed only on either transcript level (blue) or protein level (green) or on both levels (red). Red box indicates part of the volcano plot shown at higher magnification. ns: P≥0.05, *: P<0.05, **: P<0.01, ***: P<0.001, ****: P<0.0001.

Finally, we compared RNAseq and mass spectrometry data by plotting fold changes of 2,614 mRNAs against the corresponding protein variants from Uniprot database (Fig. 5H Tab. S5). This data presentation led to the identification of distinct groups, whose expression is changed on i) transcript level only (total: 20, 15 up/5 down), ii) protein level only (total: 58, 32 up/26 down) or iii) transcript and protein level (total: 10, 7 up/3 down) (Tab. S5). Interestingly, 11 out of 15 genes (73%) including Acta, Tpm2, Myl9 showing upregulation on transcript level only as well as 5 out of 32 genes (16%) including Capn2, Actn1, Cbx2 showing upregulation on protein level only have been previously identified as MRTF-SRF targets (Esnault et al., 2014; Gualdrini et al., 2016). Instead, 7/7 genes (100%), whose transcript and protein levels were both increased in CAP1-KO mice (Csrp1, Vcl, Wdr1, Flna, Cfl1, Actn3 and Fermt2), are well-established MRTF-SRF targets (Fig. 5H), suggesting that they represent core MRTF-SRF targets in E18.5 cerebral cortex. Different from MRTF-SRF targets, TCF-SRF targets were represented by only a minor fraction within upregulated genes regardless of the used approach (transcript only: 3/15 (20%), protein only: 2/32 (6%), both: 2/7 (29%)), and nearly half of them are shared with MRTF-SRF gene sets. Thus, we did not observe enrichment of TCF-SRF targets among upregulated genes in CAP1-KO mice.

In summary, RNAseq and mass spectrometry revealed a specific upregulation of MRTF-SRF targets in CAP1-KO cerebral cortex, thereby confirming our in vitro data. Moreover, these analyses led to the identification of target genes, whose expression is controlled via the CAP1-actin-MRTF-SRF pathway in neurons.

## Discussion

In the present study we identified CAP1, an ABP with largely unknown physiological functions (Rust et al., 2020; Rust and Marcello, 2022), as a crucial repressor of MRTF-SRF-mediated transcription in neurons. Our data let us propose a model in which cytosolic CAP1 promotes cytosolic MRTF retention and thereby represses MRTF-SRF-mediated transcription via an actin-dependent mechanism. It is supported by strongly increased nuclear MRTF levels and SRF activity in CAP1-KO neurons, and by our findings that both increases were fully rescued by cytosolic, but not by nuclear CAP1 and by pharmacological and genetic interventions that either increase G-actin levels or inhibit MRTF activity. Further, we identified CAP1’s HFD and CARP domain to be relevant for regulating MRTF localization and SRF activity. Specific upregulation of MRTF-SRF target genes in cerebral cortex from CAP1-KO mice, which we found by RNAseq and mass spectrometry, not only confirmed CAP1 as a crucial repressor of MRTF-SRF signaling, but also proved its *in vivo* relevance for this pathway and led to the identification of neuronal MRTF-SRF target genes.

In the present study, we showed that mutant CAP1 variants impaired in actin-binding, caused either by deletion or mutation of HFD or CARP domain (Kotila et al., 2018; Kotila et al., 2019), were not able to rescue MRTF localization or SRF activity in CAP1-KO neurons. Instead, a truncated CAP1 variant lacking C-terminal four AA residues (CAP1-Δ471-474), which have been implicated in actin regulation as well (Kotila et al., 2018), normalized both defects in CAP1-KO neurons, and we therefore concluded that these residues were dispensable for MRTF-SRF signaling. This can be explained by the fact that CAP1-Δ471-474 only displayed diminished nucleotide exchange activity on G-actin, while G-actin-binding was strongly reduced and nucleotide exchange activity was abolished upon CARP domain mutation (Kotila et al., 2018). Similar to CAP1-Δ471-474, CAP1-Δ1-35 as well as CAP1-PP1mut normalized MRTF localization and SRF activity in CAP1-KO neurons. CAP1-Δ1-35 specifically lacked OD together with two RLE motifs, which are necessary for both adenylyl cyclase binding and positive modulation of GαS-stimulated adenylyl cyclase activity in mammals (Zhang et al., 2021; Rust and Marcello, 2022), and CAP1-PP1mut is impaired in interaction with the ABP profilin or SH3 domain proteins such actin-binding protein 1 (ABP1) or tyrosine kinase ABL1 (Freeman et al., 1996; Bertling et al., 2007, Balcer, 2003 #86; Rust et al., 2020). We therefore concluded that CAP1 oligomerization, its stimulatory function towards adenylyl cyclase as well as its interaction with profilin, ABP1 or ABL1 were not important for neuronal MRTF-SRF signaling. In summary, we identified CAP1’s HFD and CARP domain to be relevant for regulating MRTF-SRF signaling, including nucleo-cytosolic MRTF shuttling. Together with altered MRTF localization upon pharmacological intervention of the actin cytoskeleton, we thereby provide strong evidences for actin-dependent nucleo-cytosolic MRTF shuttling in neurons, which has been subject of debate in the past (Kalita et al., 2006; Wickramasinghe et al., 2008; O’Sullivan et al., 2010; Stern et al., 2013).

While SRF activity was strongly increased in CAP1-KO neurons, it was unchanged upon inactivation of either cofilin1 or its close homolog ADF. We were surprised by this finding, since previous studies proved cofilin1 as a key actin regulator in neurons (Bellenchi et al., 2007; Gu et al., 2010; Rust et al., 2010; Bosch et al., 2014; Rust, 2015a; Bamburg et al., 2021). Different from Cfl1-KO or ADF-KO neurons, SRF activity was increased in ADF/Cfl1-dKO neurons, in line with previous studies that demonstrated overlapping brain functions for both ABP (Görlich et al., 2011; Flynn et al., 2012; Wolf et al., 2015; Zimmermann et al., 2015). However, SRF activity in ADF/Cfl1-dKO neurons was only moderately increased by 60%, considerably less when compared to the three- to fourfold increase in CAP1-KO neurons. In ADF/Cfl1-dKO neurons, increased SRF activity was associated with a moderate shift towards more nuclear MRTF, which was normalized upon actin depolymerization. These findings let us assume that dysregulated MRTF-SRF signaling in ADF/Cfl1-dKO neurons was caused by a primary defect in the actin cytoskeleton, as concluded for CAP1-KO neurons. However, normal SRF activity in cofilin1-KO neurons as well as the much weaker defects in MRTF localization and SRF activity in ADF/Cfl1-dKO neurons when compared to CAP1-KO neurons let us conclude that cofilin1 only plays a minor role in regulating neuronal MRTF-SRF signaling. Interestingly, previous studies placed cofilin1 downstream of SRF in neuron migration, dendritic spine morphology, axon regeneration, mitochondrial dynamics or stress resilience (Alberti et al., 2005; Beck et al., 2012; Stern et al., 2013; Zimprich et al., 2017; Nader et al., 2019). In line with these studies, we found cofilin1 among upregulated genes and proteins in cerebral cortex from CAP1-KO mice. These findings and our data let us conclude that cofilin1 acts as a crucial executor, but not as major regulator of the MRTF-SRF pathway in neurons.

In the present study, we not only solved the CAP1-dependent molecular mechanism of neuronal SRF signaling, but also proved the *in vivo* relevance of CAP1 for SRF-dependent gene expression. Specifically, a priori GSEA revealed an enrichment of established SRF targets among genes we found upregulated in cerebral cortex from CAP1-KO mice by RNAseq. The coactivators MRTF and TCF compete for SRF to introduce specificity into SRF-dependent gene expression (Olson and Nordheim, 2010), and previous landmark studies specified gene sets controlled by both coactivators (Esnault et al., 2014; Gualdrini et al., 2016). These studies proved that MRTF-SRF primarily controls expression of cytoskeletal genes to induce contractility, while TCF-SRF controls gene expression programs involved in cell signaling and proliferation. Our comparison with these gene sets revealed an enrichment of MRTF-SRF targets among upregulated genes in CAP1-KO brain. Supportively, qPCR confirmed increased expression of selected MRTF-SRF-regulated genes, but unchanged or even decreased expression of TCF-SRF-specific targets. Further, GO term analyses revealed an enrichment of genes involved in actin regulation, actin-based processes or cytoskeletal structures among upregulated genes in CAP1-KO brain, and several established MRTF-SRF targets were present among those proteins we found upregulated in cerebral cortex of CAP1-KO mice by mass spectrometry. We therefore concluded that CAP1 is specifically relevant for the MRTF-SRF pathway in cerebral cortex, in very good agreement with our data in isolated neurons.

Different from actin and key actin regulators such as cofilin1 or Arp2/3 complex subunits, CAP1 expression in fibroblasts was neither induced by serum-stimulation nor by pharmacological MRTF activation, and basal CAP1 expression was not repressed upon rise in G-actin levels (Esnault et al., 2014). CAP1 was therefore classified as a ‘no MRTF-SRF target gene’. Hence, opposite to cofilin1, CAP1 acts as a crucial regulator, but not as an executor of MRTF-SRF signaling. The fact that CAP1 acts upstream of MRTF-SRF, but itself is not controlled by this pathway let us propose an intriguing idea in which intracellular signaling cascades impinge on CAP1 to control MRTF-SRF-mediated transcription without counterbalancing changes in CAP1 expression. Intriguingly, fibroblast studies revealed induction of CAP1 expression by the ERK activator 12-O-tetradecanoyl phorbol-13-acetate (TPA), which depended on TCF-SRF activity (Gualdrini et al., 2016). Via increasing CAP1 expression, the TCF-SRF pathway may inhibit nuclear MRTF translocation and MRTF-SRF signaling, thereby inducing a proliferative, but anti-contractility gene program. It will be exciting to test in future studies whether CAP1 indeed acts as regulatory switch that controls balance between TCF-SRF and MRTF-SRF signaling. To date, several interaction partners have been identified for CAP1 or its homologs (Rust et al., 2020; Rust and Marcello, 2022), ranging from signaling molecules such as adenylyl cyclase, glycogen synthase kinase 3 (GSK3) and ABL1 to actin regulators including profilin, inverted formin 2 (INF2), cofilin1 and twinfilin (Wills et al., 2002; Balcer et al., 2003; Zhou et al., 2014; Johnston et al., 2015; A et al., 2019; Kotila et al., 2019). While our data excluded cofilin1 as a major neuronal SRF regulator and argued against important roles for CAP1’s interactions with adenylyl cyclase, profilin or ABL1 in SRF regulation, it will be exciting to test in future studies which signaling cascades act on CAP1 to control its function in balancing gene expression programs in neurons. Exemplarily, GSK3 has been reported to stimulate neuronal gene expression by SRF phosphorylation (Li et al., 2014). GSK3 also inhibits CAP1-mediated actin dynamics (Zhou et al., 2014), which may further boost its stimulatory effect on neuronal SRF activity.

KEGG analysis revealed an enrichment of genes associated with dilated cardiomyopathy (DCM) and hypertrophic cardiomyopathy (HCM) among upregulated genes in CAP1-KO brain. Interestingly, cardiac defects including DCM have been reported for systemic CAP2-KO mice and associated with human CAP2 mutations (Peche et al., 2012; Field et al., 2015; Rust and Marcello, 2022). Transcriptome analysis in CAP2-KO mice revealed an upregulation of fetal genes including SRF-regulated genes prior to cardiac defects, and SRF inhibition delayed pathology in these mice (Xiong et al., 2019). These findings and our data suggested a general role for CAPs in SRF regulation. While we found that CAP1, but not CAP2 acts as a crucial SRF regulator in the brain, CAP2, which is abundant in striated muscles and crucial not only for heart function, but also for skeletal muscle development (Kepser et al., 2019), controls SRF-dependent gene expression in the heart, in mouse fibroblasts and possibly also in skeletal muscles (Xiong et al., 2019; Kepser et al., 2021). However, different from CAP2, which has been located in both cytosol and nucleus (Peche et al., 2007), we found no evidence for nuclear CAP1 localization in neurons or brain lysates. Further, rescue experiments in CAP1-KO neurons showed that SRF was highly sensitive to cytosolic CAP1, but much less to nuclear-targeted CAP1. We therefore conclude that the MRTF-SRF pathway is primarily controlled by cytosolic CAP1.

In summary, we identified CAP1 as an important negative regulator of MRTF-SRF-mediated transcription in neurons, which promotes cytosolic MRTF retention and represses MRTF-SRF signaling by an actin-dependent mechanism that requires its HFD and CARP domain. Our study not only revealed important novel insights into regulatory mechanisms upstream of MRTF-SRF signaling in cerebral cortex, but also provide a list of MRTF-SRF-regulated neuronal genes. Further, our data propose CAP1 as a novel therapeutic avenue for the treatment of human brain disorders associated with MRTF-SRF dysregulation including Alzheimer’s disease and neuropsychiatric disorders such as autism spectrum disorders and schizophrenia (Chow et al., 2007; Holt et al., 2010; Neale et al., 2012; Luo et al., 2015; Wang et al., 2016). Considering the broad expression of CAP1, this might be relevant also for non-neuronal SRF-associated pathologies ranging from cancer to cardiovascular and metabolic diseases (Miano et al., 2010; Onuh and Qiu, 2020).

## Material and methods

### Transgenic mice

Mice were housed in the animal facility of the University of Marburg on 12-hour dark-light cycles with food and water available *ad libitum*. Treatment of mice was in accordance with the German law for conducting animal experiments and followed the guidelines for the care and use of laboratory animals of the U.S. National Institutes of Health. Generation of brain-specific CAP1-KO (CAP1^flx/flx,Nestin-Cre^) mice has been approved by RP Giessen (file references G22/2016, G13/2021) and described before (Schneider et al., 2021b). CAP2^-/-^ mice has been obtained from European Conditional Mouse Mutagenesis Program (EUCOMM). Generation of ADF^-/-^ and Cfl1^flx/flx^ mice has been described before (Bellenchi et al., 2007). Killing of mice has been approved by internal animal welfare authorities (file references AK-5-2014-Rust, AK-6-2014-Rust, AK-12-2020-Rust) and by RP Giessen (file references G22/2016, G13/2021).

### DNA constructs and oligonucleotides

pcDNA3.1-CAP1-eGFP was purchased from GenScript, where CDS of NM_007598.4 was cloned upstream of eGFP. mCherry-CAP1, MYC-CAP1 and corresponding constructs with point mutations have been described previously (Heinze et al., 2022). MYC-CAP1-WH2mut, C-terminally deleted MYC-CAP1 (Δ218-474, Δ319-474 and Δ471-474) overexpressing plasmids were generated using modified site-directed mutagenesis protocol (Liu et al., 2008). N-terminally deleted MYC-CAP1 (Δ1-35, Δ1-215 and Δ1-317) were prepared by amplification of corresponding region of CAP1 from pcDNA3.1-CAP1-eGFP and cloning between SalI and NotI restriction sites of pCMV-Myc-N. Small hairpin RNAs against CAP1 (sh1, sh4) (Zhang et al., 2013; Bertling et al., 2004), MRTFA (sh1), MRTFB (sh1) and CTR-sh were cloned into pSuper vector (OligoEngine) between BglII and HindIII restriction sites. Nuclear export signal (NES) of human PKIα (Saito et al., 2004) and nuclear localization signal (NLS) from pEF-Flag-NLS-Actin (Posern et al., 2002) was cloned into pCMV-Myc-N-CAP1 between EcoRi and SalI to generate pCMV-Myc-N-NES-CAP1 and pCMV-Myc-N-NLS-CAP1 constructs, respectively. MYC-MRTFB overexpressing constructs were created by amplification of NM_153588.3 CDS from murine brain cDNA and cloning between SalI and NotI restriction sites of pCMV-Myc-N vector. For sequences of oligonucleotides used for PCR amplification, quantitative PCR, cloning and point-mutagenesis see Tab. S6. Generation of 3D.AFOS, pRlik, pEF-Flag-MRTFA, pEF-Flag-ΔN-MRTFA, pEF-MRTFA-GFP, pCIG2-CRE-Mut-mCherry, pCIG2-CRE-mCherry, pCIG2-CRE-Mut-BFP, pCIG2-CRE-BFP, pEF-Flag-Actin, pEF-Flag-NLS-Actin and pEF-Flag-Actin-R62D have been reported before (Hill et al., 1995; Posern et al., 2002; Miralles et al., 2003; Posern et al., 2004; Kullmann et al., 2020).

### Cell culture and transfection

Primary cortical neurons from embryonic day 18 (E18) mice were prepared as previously described (Schneider et al., 2021a). In brief, cerebral cortices from the same litter were pooled-together, treated with TrypLE™ Express (Gibco) for 6 min at 37°C and dissociated with pipetting up and down 6-7 times. Cells were plated with a density of either 53,000 (for immunocytochemistry) or 132,000 (for dual-luciferase reporter assay) cells per cm^2^ on 0.1 mg/ml poly-L-lysine-coated coverslips in 24 well plates. Neurons were cultured in a humidified incubator at 37°C with 5% CO_2_ in Neurobasal medium containing 2% B27 supplement, 2 mM GlutaMax-I, 100 µg/ml streptomycin, and 100 U/ml penicillin (Gibco). Cortical neurons were transfected on DIV6/7 with 1 µg plasmid/well of 24 well plates using Lipofectamine 2000 reagent (Thermo Fischer) for 1.5 hours according to manufacturer’s protocol.

Mouse hippocampal and hypothalamic cells lines, HT-22 and mHypoE-N1, respectively, were cultured in Dulbecco’s Modified Eagle Medium containing 10% Fetal Bovine Serum, 2 mM GlutaMax-I, 100 µg/ml streptomycin, and 100 U/ml penicillin (Gibco). HT-22 and mHypoE-N1 were plated at 27,000 cells per cm^2^ in 24 well plates and next day were transfected with 1 µg plasmid/well using Lipofectamine 2000 reagent (Thermo Fischer) for 5 hours according to manufacturer’s protocol.

### Dual-luciferase reporter assay

For dual-luciferase assay, each well of 24 well plate containing DIV7 cortical neurons were transfected with 3D.AFOS (150 ng, Firefly luciferase), pRlik (150 ng, Renilla luciferase), protein overexpression constructs (usually 250 ng, pCMV-MYC-N-CAP1, pEF-Flag-Actin, pEF-MRTFA-GFP, pCMV-Myc-N-MRTFB etc..) and either pCIG2-CRE-Mut-mCherry (50 ng) or pCIG2-CRE-mCherry (50 ng). 48 hours after transfection cells were washed twice with 500 µl of 1xPBS, lysed with 100 µl of 1xPassive lysis buffer (Promega) and frozen at -20 °C for at least 1 hour. After thawing cells, 40 µl of lysate was used to measure luciferase activity with GloMax microplate reader (Promega).

### Immunocytochemistry

Cortical neurons transfected at DIV6/7 with pCIG2-CRE-Mut-BFP (150 ng) or pCIG2-CRE-BFP (150 ng) and protein overexpression constructs (usually 250 ng, pCMV-MYC-N-CAP1, pEF-MRTFA-GFP, pCMV-Myc-N-MRTFB etc..) were incubated for 3 days and at DIV9/10 were washed once with PBS, fixed in a 4% PFA/Sucrose solution for 15 min and rinsed in PBS three times. After 10 min incubation in carrier solution (CS; 0.1% gelatin, 0.3% Triton-X100 in PBS), neurons were incubated with primary antibodies (Rabbit-anti-GFP: 1:1000, Thermo Fisher Scientific, Cat. #G10362; mouse-anti-c-myc: 1:200, Thermo Fisher Scientific, Cat. #13-2500; rabbit-anti-MRTFB: 1:100, Cell signalling, Cat. #14613; rabbit-anti-DCX: 1:400, Abcam, Cat. #18723; mouse-anti-CAP1: 1:200, Abnova, Cat. # H00010487-M01) in CS for 2h. Thereafter, neurons were washed with PBS three times for 5 min and incubated with secondary antibodies (goat anti-rabbit-AlexaFluor488: 1:1000, Thermo Fisher Scientific, Cat. #A-11034; goat anti-mouse-AlexaFluor488: 1:1000, Thermo Fisher Scientific, Cat. Cat # A-11001; donkey anti-mouse-AlexaFluor546: 1:1000, Thermo Fisher Scientific, Cat. Cat # A-10036) in CS for 1 hour. After washing five times with PBS, coverslips were mounted onto microscopy slides using AquaPoly/mount (Polysciences Inc.). Neurons with nuclear and cytoplasmic localization of MRTFA or MRTFB were counted using Zeiss LSM 5 Pascal microscope, images were acquired with Leica TCS SP5 II confocal microscope setup.

### RNA-seq analysis

Total RNA from cerebral cortices of CAP1 ^flx/flx^ (n=3) and CAP1^flx/flx,Nestin-Cre^ (n=3) E18 embryos were extracted using peqGOLD TriFast reagent (PeqLab) according to manufacturer’s instructions. DNA contamination was removed by treatment with DNaseI (Qiagen) and purified with MinElute kit (Qiagen). RNA quality was determined using Experion RNA StdSens analysis kit (BioRad) by Genomics core facility, Marburg. SmallRNA libraries were constructed and sequenced by EMBL genomic core facility (Heidelberg, Germany). In brief, barcoded stranded mRNA-seq libraries were prepared from high quality total RNA samples (∼150 ng/sample) using the NEBNext Poly(A) mRNA Magnetic Isolation Module and NEBNext Ultra II Directional RNA Library Prep Kit for Illumina (New England Biolabs (NEB), Ipswich, MA, USA) implemented on the liquid handling robot Beckman i7. Obtained libraries that passed the QC step were pooled in equimolar amounts; 2.2 pM solution of this pool was loaded on the Illumina sequencer NextSeq 500 and sequenced uni-directionally, generating ∼500 million reads, each 84 bases long. Raw sequencing reads were trimmed from 3LJ-adapter (AGATCGGAAGAGC) and filtered according to quality and length using default parameters of Fastx-Toolkit for fastq data on Galaxy, a web-based genome analysis tool (https://usegalaxy.org; Afgan et al., 2018) . Reads longer than 15 nt were mapped to the mouse genome (GRCm38-mm10) with STAR RNA-aligner software (Dobin et al., 2013) and differential expression of transcripts was performed using edgeR (FDR<0.05), both of which are integrated into https://usegalaxy.org. GSEA, KEGG and GO term enrichment analysis was carried out with Perseus software version 1.6.15 (Tyanova et al., 2016). MRTF-SRF and TCF-SRF signature overrepresentation probability was calculated using hypergeometric test (https://systems.crump.ucla.edu/hypergeometric/).

### Quantitative PCR

RNA samples generated for RNA-seq experiment were treated with TURBO DNase (Thermo Fisher Scientific) according to manufacturer’s protocol. For detection of transcripts, 200ng of totalRNA sample was reverse transcribed with iScript cDNA synthesis kit (Bio-Rad) and quantitative PCR (q-PCR) was performed on the StepOnePlus Real-Time PCR System (Applied Biosystems), using iTaq SYBR Green Supermix with ROX (Bio-Rad). Each sample was measured in dub- or triplicates. qRT-PCR data were analyzed by 2^−dCt^ method, where Ct values were first normalized to an internal control (GAPDH) and then to the reference sample, which was arbitrarily set to 1. Primers used for the qPCR are provided as Tab. S6.

### Mass spectrometry

Protein lysates from CAP1 ^flx/flx^ (n=4) and CAP1 ^flx/flx, Nestin-Cre^ (n=4) cerebral cortices of E18.5 embryos were prepared by addition of 500 µl lysis buffer (0.1M Tris-HCl, pH 8.0, 0.1 M DTT, 2% SDS) to the tissue and homogenized with 20 strokes using plastic pestle in the 1.5 mL collection tube. Then lysates were passed through the 30G needle with 1 mL syringe for 5 times and incubated at 95°C for 5 min. The viscosity of the sample was reduced by sonication with Branson sonifier 450 (duty cycle 50%, 20 pulses with minimal power settings – output 10-12 %) in cold room. Non-dissolved cell debris was discarded as a pellet by 10 min centrifugation at 20°C.

Supernatants were treated with 5 mM TCEP for 15 min at 90°C. After cooling down the sample to RT, 10 mM iodoacetamide was added and the sample was incubated 30 min in the dark. Excess iodoacetamide was neutralized by the addition of an excess DTT. Samples were then further processed using a SP3 protocol adapted from previous study (Hughes et al., 2014).

4 µL of the SP3 bead-slurry was added to 25 µL of each sample (if samples had less volume, volume was adapted to 25µL by the addition of Millipore water). Subsequently 29 µL acetonitrile were added. Samples were quickly vortexed and then incubated at RT for 15 minutes. Beads were then separated using a magnetic separator. Supernatant was discarded. Beads were washed two times with 500 µL of 70% ethanol. Finally, beads were washed once with 200 µL acetonitrile. Dry beads were incubated over-night with sequencing grade modified trypsin (SERVA) in 100 µL digestion buffer (10% acetonitrile, 50 mM ammoniumbicarbonate) at 1,200 rpm at 30°C in a thermomixer.

Beads were separated using a magnetic separator. Supernatant was transferred to new 1.5 mL reaction tubes. 30 µL 2% DMSO were added to the beads and solution was incubated for 5 min in an ultrasonic bath. Subsequently beads were separated using a magnetic separator and supernatant was added to the corresponding tubes containing the supernatants. Subsequently, 30 µL of water were added to the beads, samples were vortexed and spinned down. Magnetic beads were separated again using a magnetic separator and supernatants added to the corresponding supernatants of the previous elution steps.

Finally, 10 µL of 5% TFA were added to each collection tube. Peptides were then desalted and concentrated using Chromabond C18WP spin columns (Macherey-Nagel, Part No. 730522). Finally, peptides were dissolved in 25 µL of water with 5% acetonitrile and 0.1% formic acid.

The mass spectrometric analysis of the samples was performed using a timsTOF Pro mass spectrometer (Bruker Daltonic). A nanoElute HPLC system (Bruker Daltonics), equipped with an Aurora column (25cm x 75µm) C18 RP column filled with 1.7 µm beads (IonOpticks) was connected online to the mass spectrometer. A portion of approximately 200 ng of peptides was injected directly on the separation column. Sample Loading was performed at a constant pressure of 800 bar.

Separation of the tryptic peptides was achieved at 50°C column temperature with the following gradient of water/0.1% formic acid (solvent A) and acetonitrile/0.1% formic acid (solvent B) at a flow rate of 400 nL/min: Linear increase from 2%B to 17%B within 60 minutes, followed by a linear gradient to 25%B within 30 minutes and linear increase to 37% solvent B in additional 10 minutes. Finally, B was increased to 95% within 10 minutes and hold for additional 10 minutes. The built-in “DDA PASEF-standard_1.1sec_cycletime” method developed by Bruker Daltonics was used for mass spectrometric measurement.

Data analysis was performed using MaxQuant (version 1.6.17.0) with Andromeda search engine against the Uniprot database. Peptides with minimum of seven amino-acid lengths were used and FDR was set to 1% at the peptide and protein level. Protein identification required at least one razor peptide per protein group and label free quantification (LFQ) algorithm was applied. Bioinformatics analysis was performed with Perseus software (1.6.15.0) using LFQ intensity values. Values were normalized by subtraction of median. Statistical analysis of differential expression was estimated using ANOVA (s0=0, permutation-based FDR= 0.05, number of randomizations=250).

### Nuclear fractionation

For nuclear fractionation, one half of cerebral cortices from postnatal day 12 (P12) mouse brain was homogenized in Dounce potter with 5 strokes in 1 mL ice-cold hypotonic homogenation buffer (HHB: 10 mM KCl, 1.5 mM MgCl_2_, 1 mM Na-EDTA, 1 mM Na-EGTA, 10 mM Tris-HCl pH 7.4, 1 mM dithiothreitol, 1x Proteinase inhibitors (Complete, Roche)). Lysate was incubated on ice for 30 min, then 1 mL of HHB with 0.2% Igepal CA-630 was added and further homogenized with 10 stokes using Dounce potter. Cortical lysate was centrifuged at 720xg for 5 min at 4°C and cytoplasmic (supernatant) fraction was collected. Nuclear fraction (pellet) was washed with isotonic homogenation buffer (HHB + 0.25 M sucrose) and passed through the 26G needle twice to clear from contaminants of cytoplasmic fraction.

### Immunoblots

Cortices of E18.5 mice were homogenized in 500 µl RIPA buffer and cultured cells were lysed in 100µl RIPA buffer containing 50 mM Tris HCl pH 7.5, 150 mM NaCl, 0.5% Igepal-CA630, 0.1% SDS and protease inhibitor (Complete, Roche). After centrifugation at 14,000 rpm for 10 min at 4°C, samples were boiled for 5 min at 95°C in Laemmli buffer including 6% DTT. Equal protein amounts were separated by SDS PAGE and blotted onto a polyvinylidene difluoride membrane (Merck) by using a Wet/Tank Blotting System (Biorad). Membranes were blocked in Tris-buffered saline (TBS) containing 5% milk powder and 0.1% Tween-20 for 1 hour and afterwards incubated with primary antibodies in blocking solution over night at 4°C. As secondary antibodies, horseradish peroxidase (HRP)-conjugated antibodies (1:20,000, Thermo Fisher Scientific) or fluorescent-conjugated antibodies (1:15,000, Li-Cor Bioscience) were used and detected by chemiluminescence with ECL Plus Western Blot Detection System (GE Healthcare) or by fluorescence with Li-Cor Odyssey imaging system. Following primary antibodies were used: mouse anti-CAP1 (1:1,000, Abnova, Cat. # H00010487-M01), mouse anti-GAPDH (1:1,000, R&D System, Cat. #MAB5718), rabbit anti-vinculin (1:500, Sigma-Aldrich, Cat. #V4139), rabbit anti-twinfilin1 (1:1000, Proteintech, Cat. #11732-1-AP), mouse anti-ADF (1:750, Sigma-Aldrich, Cat. #D8815), rabbit anti-cofilin (1:1000, Bellenchi et al., 2007) and rabbit anti-HDAC2 (1:1000, Cell signaling, Cat. #2540s).

### Statistical analysis

Experiments are reported as mean ± standard error of the mean (SEM) and (if not otherwise stated) based on three independent biological replicates. Statistical tests were done using GraphPad Prizm 9.5. For comparing only two groups, student’s t-test; more than two groups, one-way Anova with Sidak’s or Tukey’s post-hoc comparison was applied. Comparison of nuclear and cytoplasmic enriched neuron groups was performed with χ2-test.

## Acknowledgments

We thank the EUCOMM consortium for providing CAP2^-/-^ mice, Dr. Walter Witke (Bonn, Germany) for providing ADF^-/-^ mice, Cfl1^flx/flx^ mice and cofilin1 constructs, Dr. Richard Treisman for providing 3D.AFOS, pRlik and MRTFA constructs, Dr. David Solecki for providing Cre and Cre-mut constructs, Dr. Guido Posern for actin constructs and Eva Becker, Flynn Kubeja and Renate Gondrum for excellent technical support. This work was supported by research grants from Fondazione Cariplo (2018-0511) and from Deutsche Forschungsgemeinschaft (DFG, German Research Foundation; RU 1232/7-1, RU 1232/10-1) to MBR.

## Author contributions

Experiments were performed by SK, KW, AH, NT, UL and DC, experiments were designed and results were discussed by SK, KW, AH, UL and MBR, manuscript was written by MBR and SK.

