## Supplementary figures and images for "Cyclase-associated protein 1 (CAP1) represses MRTF-SRF-dependent gene expression in the mouse cerebral cortex"

### Figure S1

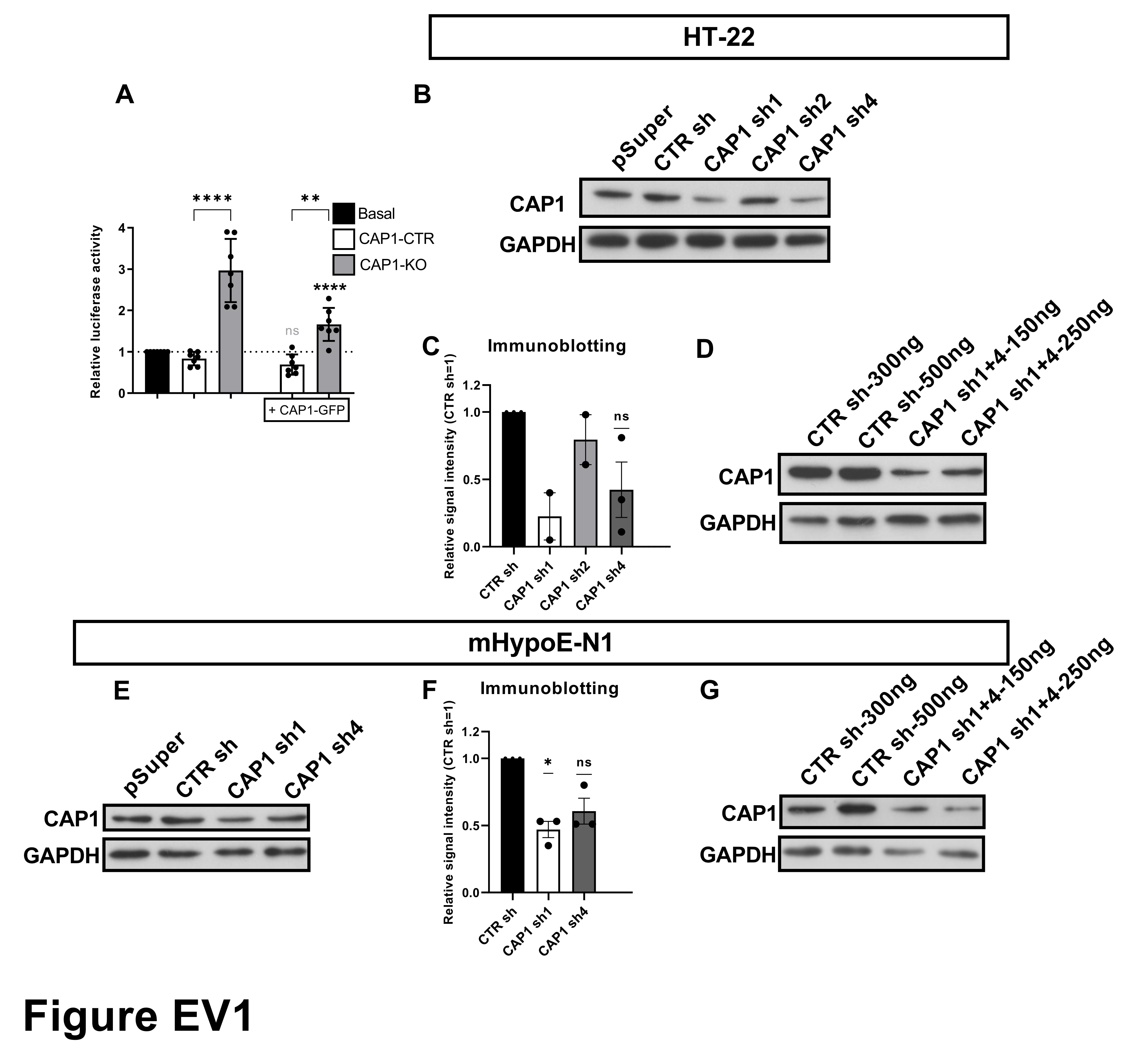

### Figure S2

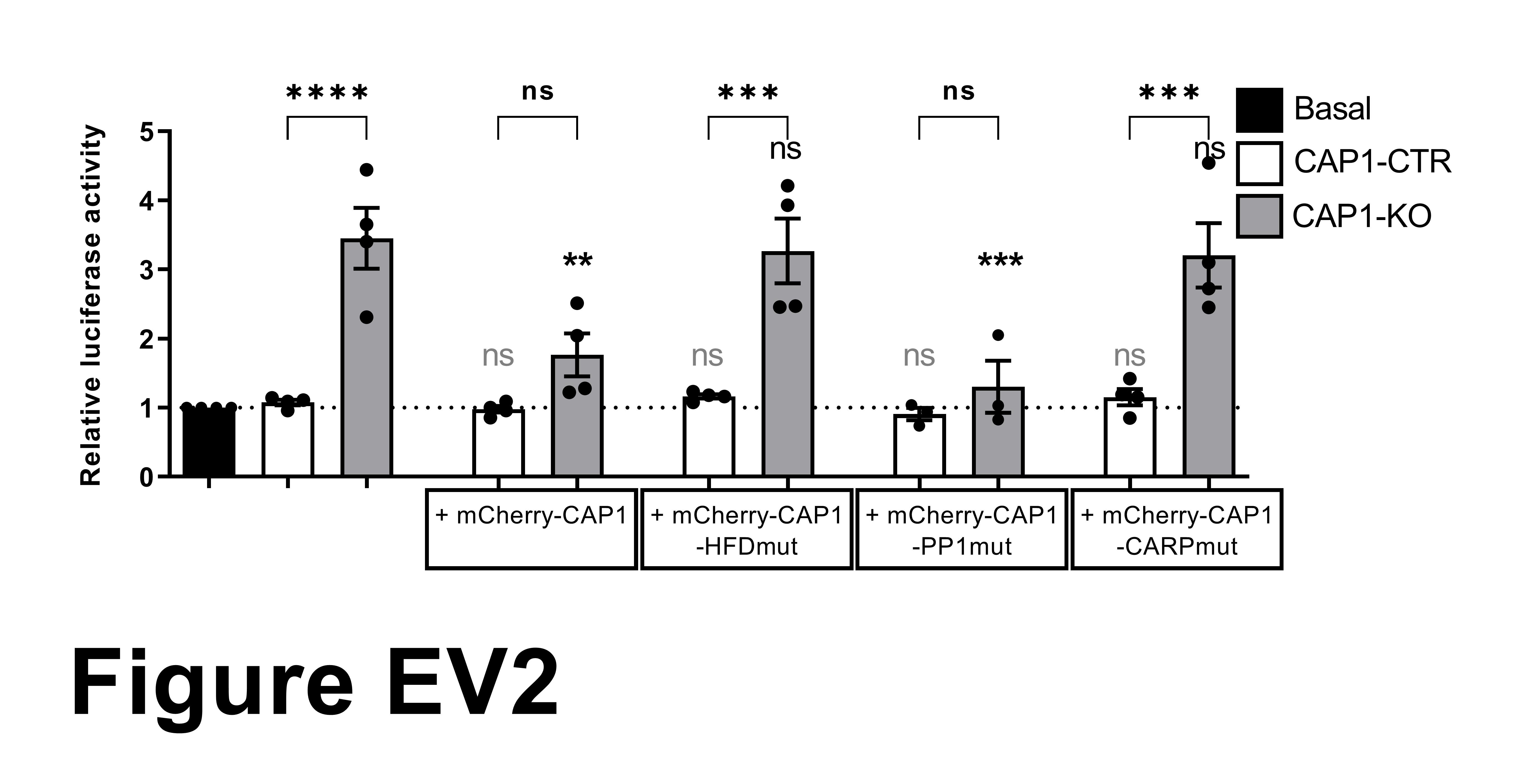

### Figure S3

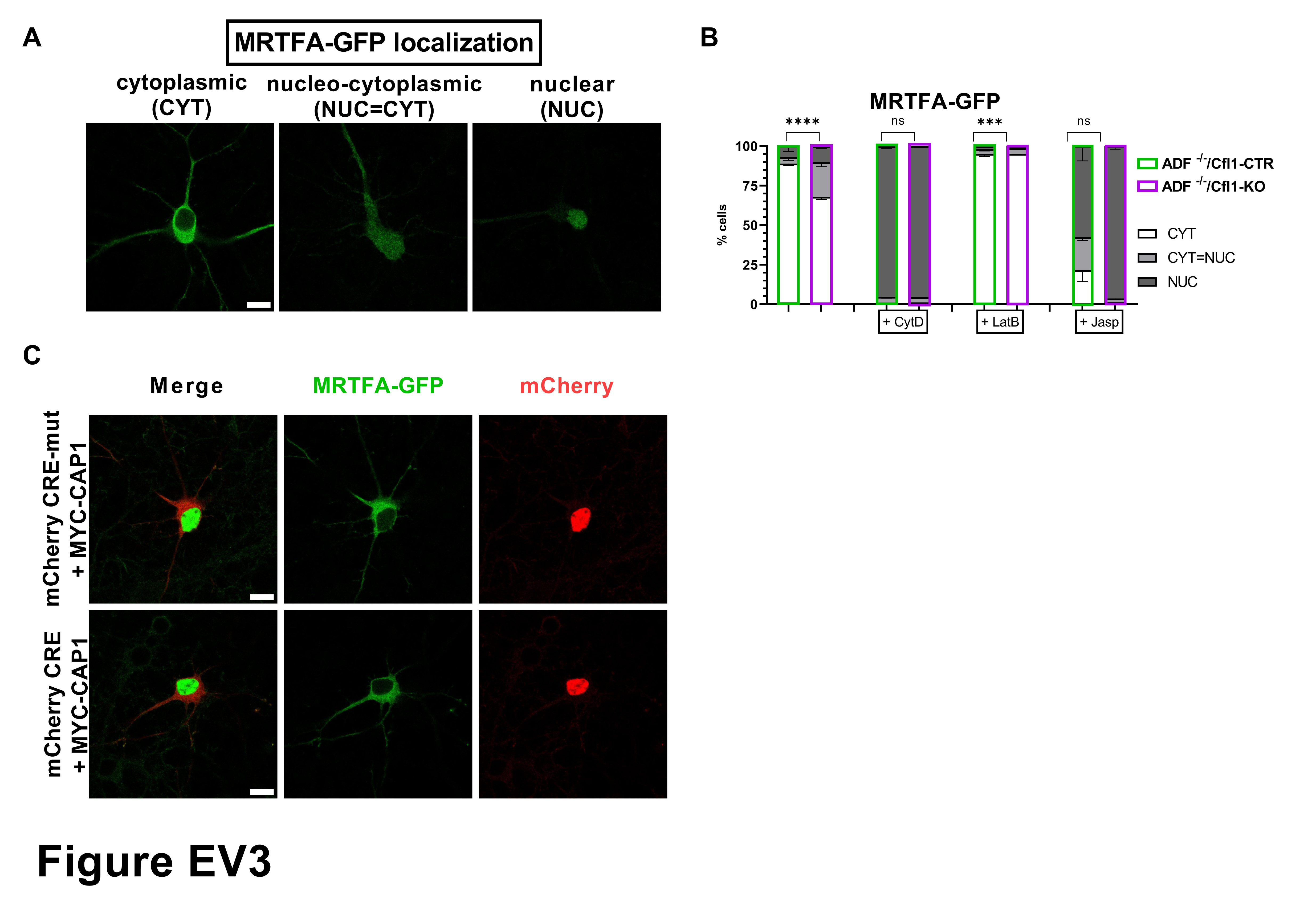
